# SpaRx: Elucidate single-cell spatial heterogeneity of drug responses for personalized treatment

**DOI:** 10.1101/2023.08.03.551911

**Authors:** Ziyang Tang, Xiang Liu, Zuotian Li, Tonglin Zhang, Baijian Yang, Jing Su, Qianqian Song

**Affiliations:** Department of Computer and Information Technology, Purdue University, Indiana, USA; Department of Biostatistics and Health Data Science, Indiana University School of Medicine, Indiana, USA; Department of Computer Graphics Technology, Purdue University, Indiana, USA; Department of Statistics, Purdue University, Indiana, USA; Center for Cancer Genomics and Precision Oncology, Atrium Health Wake Forest Baptist Comprehensive Cancer Center, North Carolina, USA; Department of Cancer Biology, Wake Forest University School of Medicine, North Carolina, USA

**Keywords:** Graph transformer, adversarial learning, spatial cellular drug response, single-cell spatial transcriptomics

## Abstract

Spatial cellular heterogeneity contributes to differential drug responses in a tumor lesion and potential therapeutic resistance. Recent emerging spatial technologies such as CosMx SMI, MERSCOPE, and Xenium delineate the spatial gene expression patterns at the single cell resolution. This provides unprecedented opportunities to identify spatially localized cellular resistance and to optimize the treatment for individual patients. In this work, we present a graph-based domain adaptation model, SpaRx, to reveal the heterogeneity of spatial cellular response to drugs. SpaRx transfers the knowledge from pharmacogenomics profiles to single-cell spatial transcriptomics data, through hybrid learning with dynamic adversarial adaption. Comprehensive benchmarking demonstrates the superior and robust performance of SpaRx at different dropout rates, noise levels, and transcriptomics coverage. Further application of SpaRx to the state-of-art single-cell spatial transcriptomics data reveals that tumor cells in different locations of a tumor lesion present heterogenous sensitivity or resistance to drugs. Moreover, resistant tumor cells interact with themselves or the surrounding constituents to form an ecosystem for drug resistance. Collectively, SpaRx characterizes the spatial therapeutic variability, unveils the molecular mechanisms underpinning drug resistance, and identifies personalized drug targets and effective drug combinations.

**Key Points:**

- We have developed a novel graph-based domain adaption model named SpaRx, to reveal the heterogeneity of spatial cellular response to different types of drugs, which bridges the gap between pharmacogenomics knowledgebase and single-cell spatial transcriptomics data.
- SpaRx is developed tailored for single-cell spatial transcriptomics data and is provided available as a ready-to-use open-source software, which demonstrates high accuracy and robust performance.
- SpaRx uncovers that tumor cells located in different areas within tumor lesion exhibit varying levels of sensitivity or resistance to drugs. Moreover, SpaRx reveals that tumor cells interact with themselves and the surrounding microenvironment to form an ecosystem capable of drug resistance.

## INTRODUCTION

Understanding how different cells spatially localize and communicate in their microenvironment are critical to personalized treatment^1,2^. For instance, tumor cells of one patient are spatially heterogenous that present differential responses to treatment. For those tumor cells, their protective microenvironment and intercellular communications contribute to therapeutic failure and disease relapse^3,4^. Therefore, in order to precisely treat a patient, it is crucial to identify that, inside its tumor lesion, which tumor cells are resistant to a candidate drug and what cell-cell communications are responsible for such tumor cell resistance. Unfortunately, these issues are not satisfactorily addressed, largely due to the lack of biotechnologies to accurately delineate the spatial heterogeneity of cells within tissues.

Recently, the emerging single-cell spatial transcriptomics (SCST) technologies, such as NanoString’s CosMx™ SMI^5^ and Vizgen’s MERSCOPE^6^, hold the promise to unravel the spatial tissue architectures at subcellular level and to further our understanding of the underlying functional mechanisms of tumor metastasis^7^ and drug resistance^8^. The SCST technologies provide the spatial locations of cells as well as their gene expression patterns, which offers a unique opportunity to investigate the therapeutic heterogeneity in the tumor microenvironment. Moreover, the existing pharmacogenomics databases, including the Cancer Cell Line Encyclopedia (CCLE)^9^ and the Genomics of Drugs Sensitivity in Cancer (GDSC)^10,11^, provide valuable references relating gene expression patterns to drug response and treatment efficacy. The integration of these existing resources, along with SCST data, presents unprecedented opportunities to elucidate how individual cells within complex tissues will differentially respond to drugs, thus meeting the needs of personalized treatment.

Given the available data from drug screening cell lines, several studies have explored the connection between cell line profiles and single-cell RNA sequencing (scRNA-seq) data to investigate drug response at the single-cell level. For instance, Gambardella et.al.^12^ proposed a method called DREEP, which utilizes scRNA-seq data to predict the drug sensitivity of individual cells. They discovered that cells exhibiting transcriptional heterogeneity displayed varying degrees of drug sensitivity. Chen et.al.^13^ developed a deep transfer learning framework, scDEAL, which integrates large-scale bulk cell-line data with scRNA-seq data to predict the response of single cells to cancer drugs. Similarly, Zheng et.al.^14^ introduced SCAD, an adversarial discriminative domain adaptation framework that leverages scRNA-seq data from the GDSC database to identify drug sensitivities. However, these methods are not directly applicable to SCST data, as they do not take into account the spatial cell locations. To address this limitation, graph-based domain adaptation models have emerged as promising solutions for uncovering the spatial cellular response to drugs. Such graph-based model aggregates information not only from individual cells but also from their spatial neighbors, enabling a more comprehensive understanding of drug response within a spatial context.

Herein, we present SpaRx, a graph-based domain adaptation model, to reveal spatial therapeutic complexity with distinct drug responses, through leveraging the high-throughput pharmacological profiles and the SCST datasets. SpaRx is able to identify the cellular drug responses within a complex tissue, the spatial surrounding microenvironment of resistant cells, and the cell-cell communications involved in drug resistance. SpaRx can accurately transfer drug response predictors trained on a source domain (e.g., cell lines) to a target domain (e.g., spatial tumor cells). Our model explicitly considers the fundamental differences between source and target domain rather than modeling these differences as technical batch effects only. SpaRx will facilitate mechanistic studies for overcoming drug resistance and advance therapeutic research for precision medicine. It also holds promises to prioritize candidate drugs for individual patients, provide therapeutic guides for synergistic drug combinations, and repurpose anti-cancer drugs for other diseases such as Alzheimer’s Disease.

## RESULTS

### Overview of SpaRx

SpaRx employs a novel domain adaption strategy to transfer the knowledge of drug responses from large-scale drug screening databases (source domain) to predict the drug-sensitivity of cells in SCST data (target domain). We hypothesize that there are domain-invariant relations between molecular profiles and drug responses. SpaRx is built to learn such transferable knowledge of drug responses across the source and the target domain through end-to-end adversarial training (**Fig. 1a**). As shown in (**Fig. 1b**), SpaRx consists of a feature extractor to project molecular profiles to a latent space, a drug response predictor to predict cellular sensitivity to a drug according to the latent representation of gene expression patterns, and three domain discriminators to distinguish the source domain from the target domain. A hybrid learning strategy is used, with an adversarial learning procedure (*L*_*t*_) involving the feature extractor and the domain discriminators to learn domain-invariant molecular features, and a supervised procedure (*L*_*y*_) involving the feature extractor and the predictor to learn molecular features that are responsible for drug responses. With this hybrid learning strategy, SpaRx can transfer the domain-invariant information from the source knowledgebase to predict the drug responses of individual cells in SCST data.

**Fig. 1:**
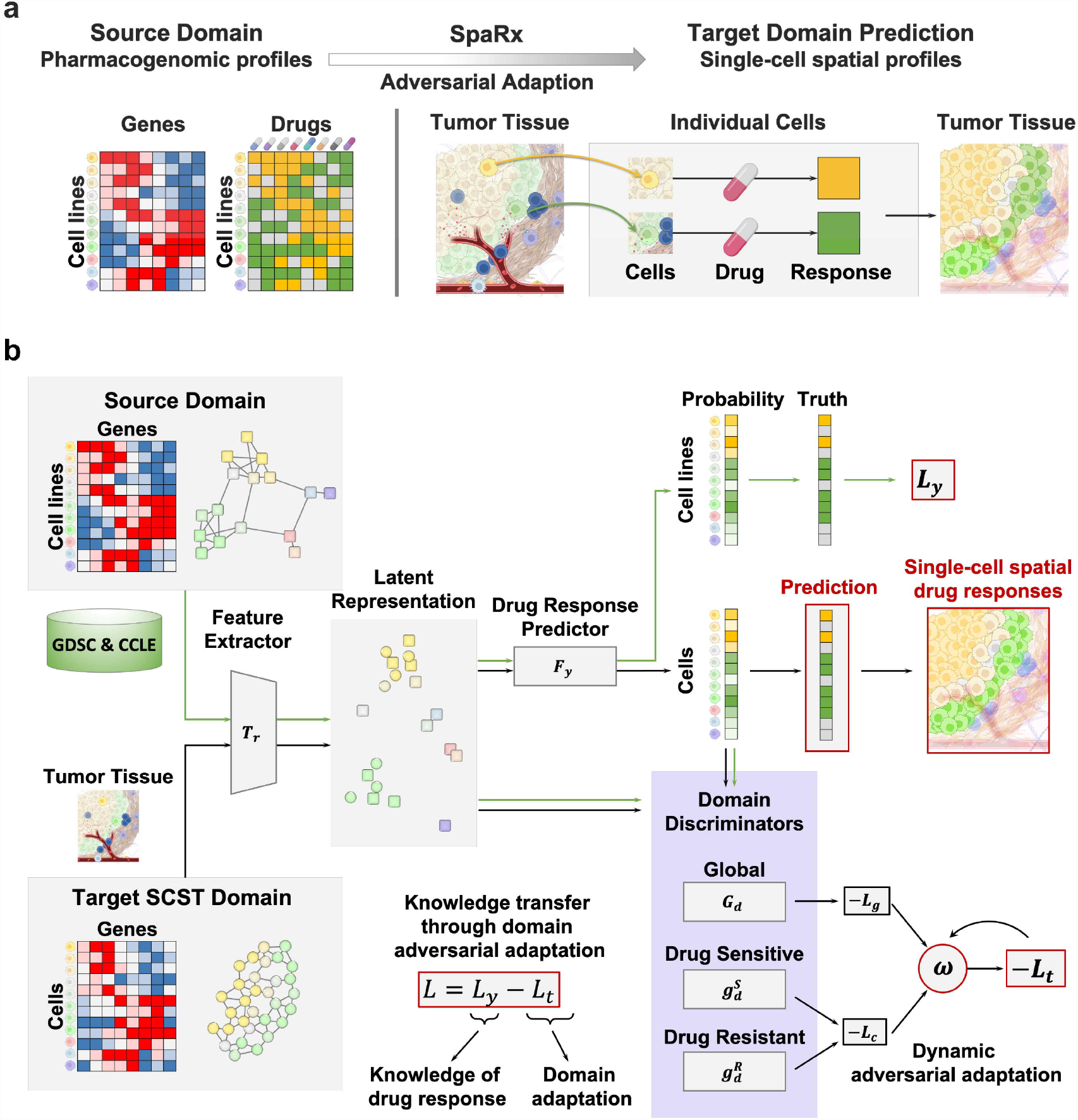
Schematic overview of the SpaRx method. **a**, Concept design of the SpaRx model. With the pharmacogenomic profiles serving as the source data, SpaRx enables to predict the cells’ response to drugs in single-cell spatial transcriptomics data via graph-based adversarial adaptation. **b**, SpaRx comprises three components (feature extractors, drug-response predictor, and discriminators) to leverage source domain and predict cellular response in target domain.

In the adversarial learning procedure, SpaRx includes dynamic adversarial adaption learning to balance the learning of global and drug-specific domain-invariant gene expression patterns. To achieve this, three discriminators are used: a global discriminator that distinguishes the source domain from the target domain for all cells or cell lines, and two drug-specific discriminators to distinguish the source domain from the target domain under each category, drug-sensitive and drug-resistant. A dynamic learnable factor (ω) is used to balance the contribution of these discriminators. Trained through the predictor-based loss that captures the knowledge of drug response, as well as the domain adaptation loss regarding global and drug-specific distributions, SpaRx is able to predict cellular sensitivity to drugs in SCST data. In this way, SpaRx successfully translates preclinical knowledge into cell-level drug response in SCST data, which facilitates deep insights into the underlying mechanisms of drug resistance and advances therapeutic effectiveness.

### SpaRx demonstrates accurate predictions of drug response

As SpaRx is the first method to predict drug response in SCST data, here we benchmark it against four deep learning (DL) models including SpaRx-GAT, SpaRx-GCN, scDEAL^13^, and SCAD^14^, as well as four machine learning (ML) methods including SVM, RF, LightGBM, and XGBoost (see **Materials and Methods**). Among them, SpaRx, SpaRx-GAT, and SpaRx-GCN share similar model architecture but use graph transformer^15^, GAT^16^, and GCN^17^ as the feature extractor. For benchmarking datasets, we randomly select a proportion (*p*) of cell lines from pharmacological database as the source domain. The remaining cell lines (1-*p*) are used to synthesize the single-cell gene expression data in target domain, with cellular complexity and drug response generated (see **Materials and Methods**). Benchmarking performance is evaluated by the accuracy of predicted drug response in target domain.

First, we randomly select 30% of cell lines (*p* = 30%) for the source domain data, and the other 70% for synthesizing the target domain data. The performance of SpaRx and other methods for predicting drug responses in the target domain are measured using the F1 score. SpaRx consistently demonstrates better performance across 80 drugs than the other methods including SpaRx-GAT (**Fig. 2a**), SpaRx-GCN (**Fig. 2b**), RF (**Fig. 2c**), and SVM (**Fig. 2d**), as well as SCAD, scDEAL, LightGBM, and XGBoost (**Supplementary Fig. 1a**). SpaRx achieves the highest accuracy (median F1 = 0.938, **Fig. 2e**), which is significantly higher than other DL models (median F1 of SpaRx-GAT: 0.787; SpaRx-GCN: 0.751; SCAD: 0.856, scDEAL: 0.669) and ML methods (RF: 0.628; SVM: 0.564; LightGBM, 0.576; XGBoost: 0.588). Meanwhile, SpaRx demonstrates particularly higher accuracy relative to SpaRx-GAT and SpaRx-GCN for certain drugs. For example, SpaRx shows noticeably higher F1 scores than SpaRx-GAT (F-1 scores, 0.923 vs 0.683) based on a hormone therapy drug tamoxifen, and higher than SpaRx-GCN (F-1 scores, 0.921 vs 0.445) for the other kinase inhibitor drug alisertib.

**Fig. 2:**
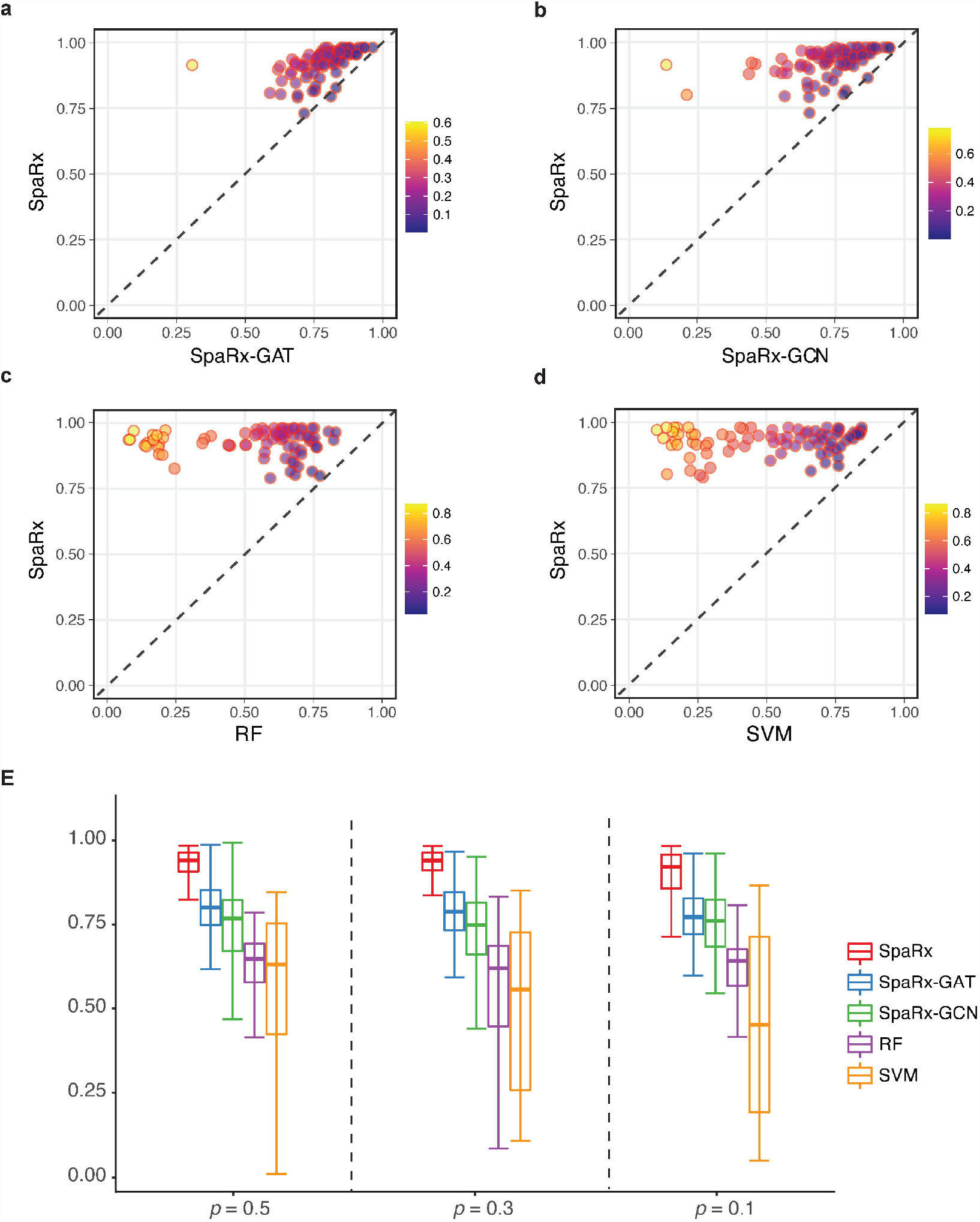
SpaRx accurately identifies drug response in targe domain. **a**, Performance of SpaRx and other methods (SpaRx-GAT, SpaRx-GCN, RF, and SVM) are measured by the F1 scores across 80 different drug compounds. Each point represents the F1 score of SpaRx versus an alternative method on one type of drug. **b**, Boxplot of F1 scores based on different sizes of source data.

Moreover, we evaluate the performance of SpaRx in the settings of different *p* (50%, 30%, 10%) based on the F1 score. Across these different settings, SpaRx (median F1 = 0.934, **Fig. 2b, Supplementary Fig. 1b**) is consistently superior to other DL models (median F1; SpaRx-GAT: 0.784; SpaRx-GCN: 0.765; SCAD: 0.873; scDEAL: 0.669), and ML methods (median F1; RF: 0.638; SVM: 0.581; LightGBM: 0.585, XGBoost: 0.604). For example, for a commonly used liver cancer drug mitoxantrone, SpaRx shows superior performance (median F1 = 0.957) than other DL models (SpaRx-GAT: 0.844; SpaRx-GCN: 0.657; SCAD: 0.678; scDEAL: 0.662). In addition to F1 score, metrics including AUROC, AUPR, precision, and recall (**Supplementary Figs 2-5, Supplementary File 1**), further demonstrate that SpaRx not only presents superior performance with different sizes of source data, but also achieves accurate response predictions for different types of drugs.

### SpaRx accurately predicts drug response in different scenarios

We further evaluate the performance of SpaRx in the scenarios of different noise levels, dropout rates, and numbers of genes in the source and the target data (see **Materials and Methods**). Benchmarking methods including four DL models (SpaRx-GAT, SpaRx-GCN, SCAD, scDEAL) and four ML models (RF, SVM, LightGBM, XGBoost).

**Fig. 3a** shows the F1 scores achieved by different methods for each drug at the noise level of 1. SpaRx achieves higher F1 scores than SpaRx-GAT and SpaRx-GCN (median F1; 0.938, 0.787, 0.751), and also performs significantly better than SCAD and scDEAL (median F1: 0.856, 0.669, **Supplementary Fig. 1b**). Moreover, when the noise level increases (noise level = 1, 1.5, 2), SpaRx maintains accurate predictions with median F1 as 0.938, 0.921, and 0.893, respectively (**Fig. 3b**). In contrast, the other methods, such as SpaRx-GAT and SpaRx-GCN, are affected by the increased noise in source and target data, indicating these methods are more likely to be undermined by data noise. The other metrics including AUROC, AUPR, precision, and recall (**Supplementary Figs 2-5, Supplementary File 1**) supports that SpaRx is robust to data noise in real applications.

**Fig. 3:**
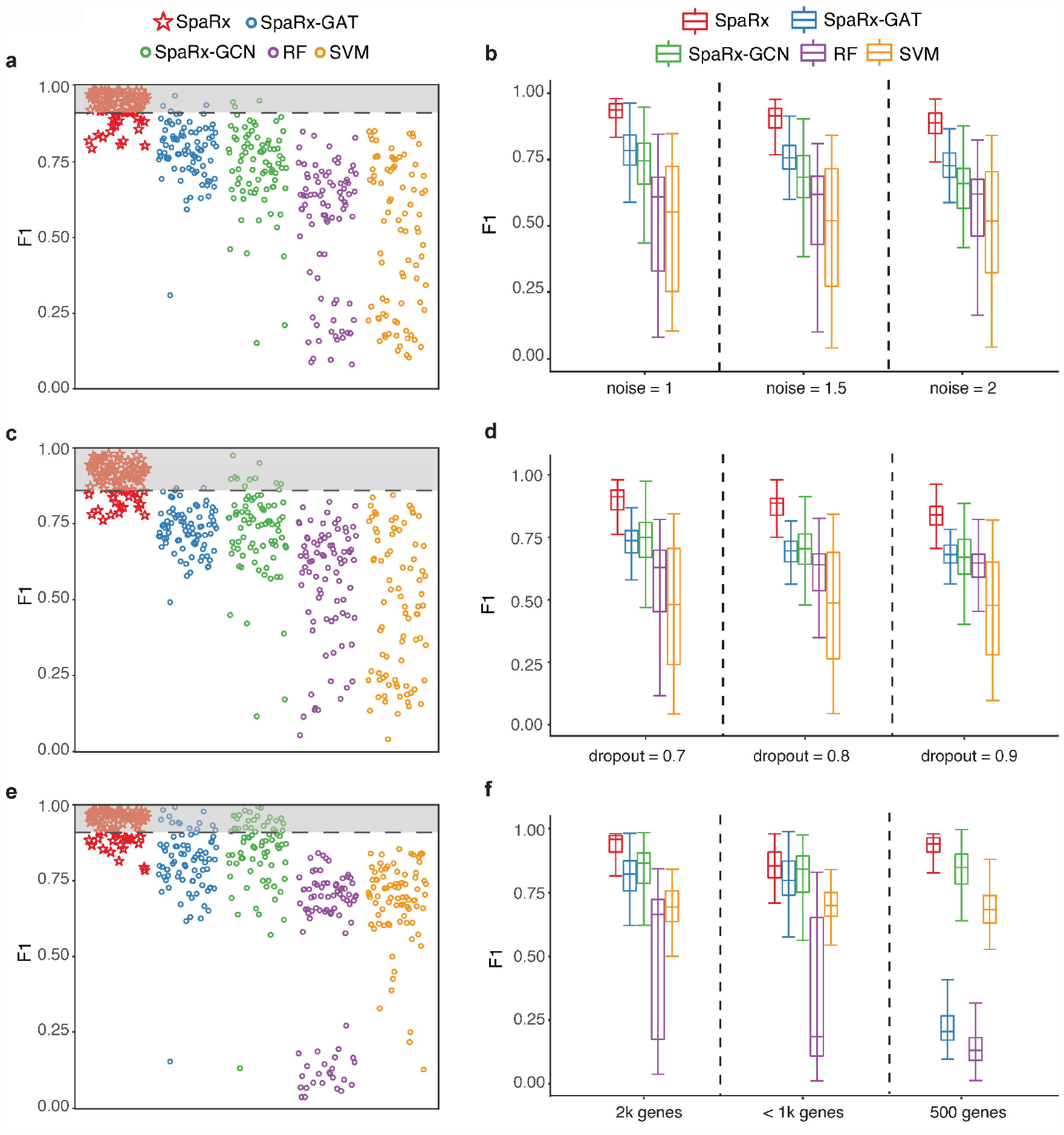
Performance evaluation in different benchmarking scenarios. **a**, Accuracy of identifying ground truth labels for the scenario with extra noise (noise level = 1) in source and target domain. Dashed lines refer to 15% percentile of SpaRx’s F1 scores. **b**, Boxplot of F1 scores over three scenarios with extra noise (noise level = 1, 1.5, 2) in source and target data. **c**, Accuracy of identifying ground truth labels for the scenario with extensive dropouts (dropout rate = 70%) in source and target domain. **d**, Boxplot of F1 scores over three scenarios with different dropout levels (dropout rate = 70%, 80%, 90%) in source and target domain. **e**, Accuracy of identifying ground truth labels for the scenario with limited number of genes (number of genes = 2k) in source and target domain. **f**, Boxplot of F1 scores over three scenarios with limited number of genes (number of genes = 2k, 853, 500) in source and target domain.

In addition, we evaluate the performance of SpaRx with different dropout rates. When the dropout rate is 70%, SpaRx remains more accurate than SpaRx-GAT and SpaRx-GCN (median F1: 0.916, 0.738, 0.751; **Fig. 3c**), as well as SCAD and scDEAL (median F1: 0.846, 0.650; **Supplementary Fig. 1b**). When the dropout rate increases (**Fig. 3d, Supplementary Fig. 1b**), SpaRx is still superior to the other DL methods (median F1; SpaRx: 0.882; SpaRx-GAT: 0.709; SpaRx-GCN: 0.707; SCAD: 0.833; scDEAL: 0.620) and ML models (median F1; RF: 0.675, SVM: 0.709, LighGBM: 0.641, XGBoost: 0.615). Other metrics including AUROC, AUPR, precision, and recall (**Supplementary Figs 2-5, Supplementary File 1**) demonstrate that SpaRx provides accurate predictions at different dropout levels.

Finally, we evaluate the performance of SpaRx with reduced number of genes in source and target data. Based on only 2,000 genes, SpaRx remains superior to SpaRx-GAT and SpaRx-GCN (median F1; 0.960, 0.823, 0.883; **Fig. 3e**) as well as SCAD and scDEAL (median F1: 0.853, 0.669; **Supplementary Fig. 1b**). With the number of genes decreasing to around 1k genes and 500 genes captured by NanoString CosMx and Vizgen MERSCOPE respectively, SpaRx maintains much more reliable performance than the other methods (median F1; SpaRx: 0.930; SpaRx-GAT: 0.753; SpaRx-GCN: 0.866; SCAD: 0.754; scDEAL: 0.595; **Fig. 3f and Supplementary Fig. 1b**). The other metrics including AUROC, AUPR, precision, and recall (**Supplementary Figs 2-5, Supplementary File 1**) shows that SpaRx outperforms benchmarking methods.

Collectively, these results demonstrate that SpaRx achieves superior predictions in different scenarios, even when the target data has extra noises, high dropout rates, and limited number of genes. These evaluations demonstrate the effectiveness of SpaRx in transferring drug-related intrinsic information across different biological domains, which enable to predict cellular drug response in single-cell spatial transcriptomics data.

### SpaRx reveals the spatial cellular heterogeneity of drug response in lung cancer

To reveal spatial cell variability in drug response, we first apply SpaRx to the NanoString CosMx lung cancer SCST data with different cell types on eight Filed Of View (FOV)^5^ (**Fig. 4a**). The zoomed-in image on the right shows that tumor cells (colored in light blue) are infiltrated with immune cells such as macrophage (colored in orange) and B cells (colored in green). Based on this tissue slice, we apply SpaRx to predict the tumor cells’ response to a typical lung cancer drug, cisplatin, for which the mechanism of action is to cause DNA damage in cancer cells, blocking cell division and leading to apoptotic cell death. As in **Fig. 4b**, SpaRx uncovers tumor cells’ response to cisplatin, which exhibits strong heterogeneity of sensitivity and resistance. Interestingly, in contrast to the agminated resistant cells, the zoom-in FOV presents a scattered pattern of sensitive cells.

**Fig. 4:**
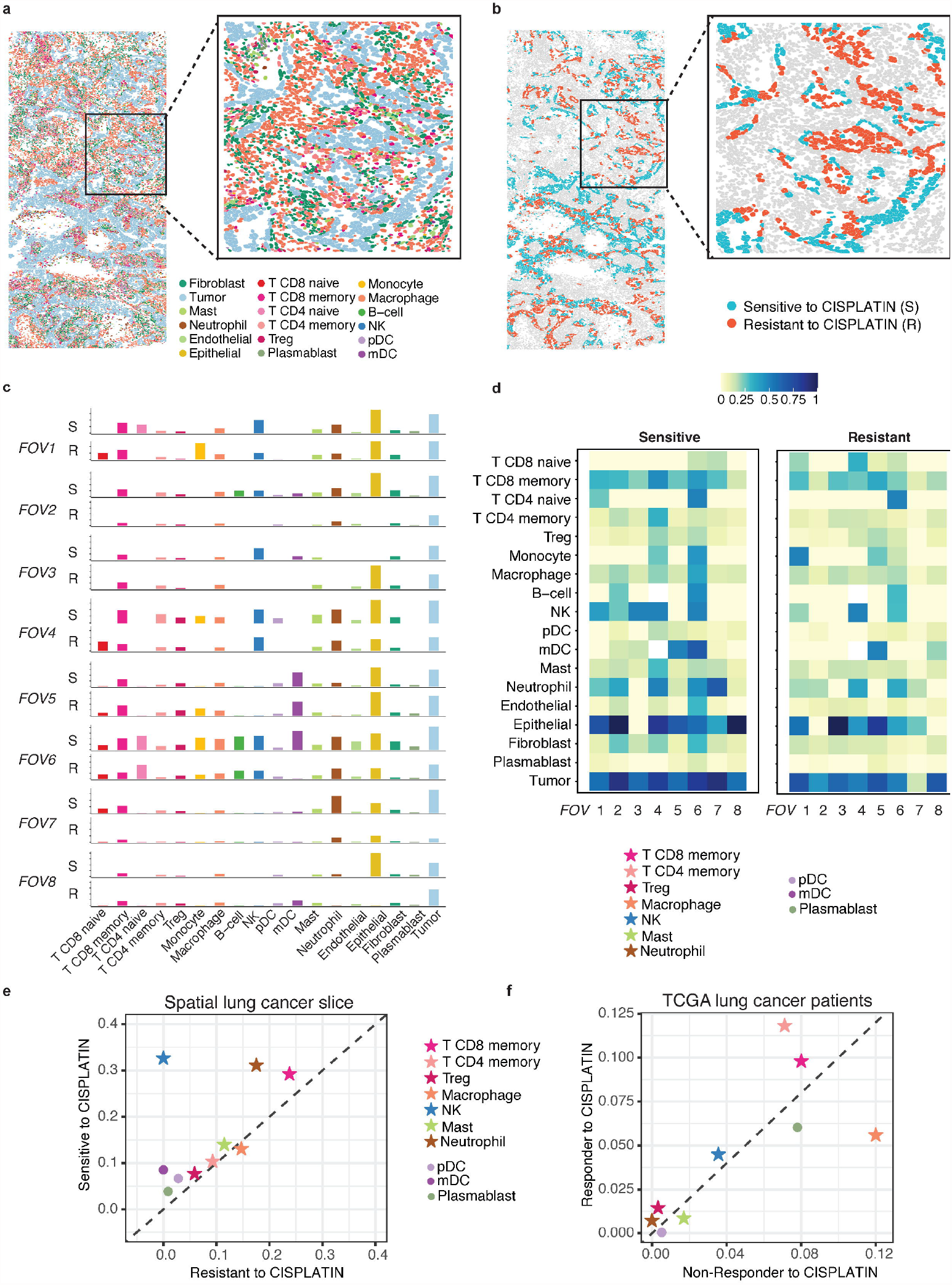
Spatial heterogeneity of cellular response to cisplatin. **a**, Spatial visualization of single-cell spatial data from lung cancer tissue. Eight Field Of Views (FOVs) are included. Different colors denote different types of cells. **b**, Spatial visualization of cellular response to cisplatin. Red and Cyan colors denote resistant (R) and sensitive (S) tumor cells to cisplatin. **c**, Surrounding microenvironment of resistant (R) and sensitive (S) cells in each FOV. **d**, Heatmap quantitatively delineates the constituents around sensitive and resistant cells. **e**, Summarization across eight FOVs for the cell type distributions surrounding resistant and sensitive tumor cells to cisplatin. **f**, Characterization of the infiltrated cell types in responders and non-responders to cisplatin in TCGA lung cancer patient samples.

Given that those tumor cells respond differentially to cisplatin, we further interrogate if the surrounding microenvironment of resistant tumor cells is different from that of sensitive ones. As shown in **Fig. 4c**, for each FOV, the spatial distributions (i.e., proportions) of cell types adjacent to resistant and sensitive cells are different. Such differences also appear to be distinct across different FOVs. Noteworthy, across FOV 1-3 (**Fig. 4c**), CD8 memory T cells, B cells, and natural killer (NK) cells are consistently reduced in the surroundings of resistant cells. Heatmap in **Fig. 4d** further quantitatively delineates the distinctive microenvironment between sensitive and resistant cells, where CD4 and CD8 memory T cells are less infiltrated in the surrounding of resistant cells. Moreover, fewer B cells, dendritic cells (DC), and NK cells are present in the microenvironment of resistant cells. Further averaging of the surrounding cell type proportions across the eight FOVs (**Fig. 4e**) shows that most cell types except macrophages are more abundant in the microenvironment of sensitive cells. Similar patterns are also observed in the TCGA lung cancer patients receiving cisplatin treatment (**Fig. 4f**). Specifically, after bulk RNA-seq decomposition by CIBERSORT^18^, CD4 and CD8 memory T cells, B cells, and NK cells are shown to be more prevalent in responders than non-responders to cisplatin. These results indicate that the surrounding microenvironment may be relevant to or modulate the tumor cells’ responses to cisplatin.

### Spatial cellular crosstalk mediates drug resistance

Given the distinctive microenvironment surrounding sensitive and resistant tumor cells, further characterization of cell-cell communications can explain how neighboring cells modulate the differential responses of tumor cells to cisplatin. Using spaCI^19^, a cell-cell communication tool specifically designed for SCST data, we infer the ligand-receptor (L–R) interactions involving tumor cells in the zoom-in FOV (**Fig. 4a**). The aggerated L–R interactions that occur between tumor cells and adjacent cells are shown in the chord diagram (**Fig. 5a**), with the chord width indicating the interaction strength. Of note, we observe that macrophage, fibroblast, and CD4 and CD8 memory T cells interact strongly with both sensitive cells and resistant tumor cells. NK cells uniquely crosstalk with sensitive but not resistant tumor cells.

**Fig. 5:**
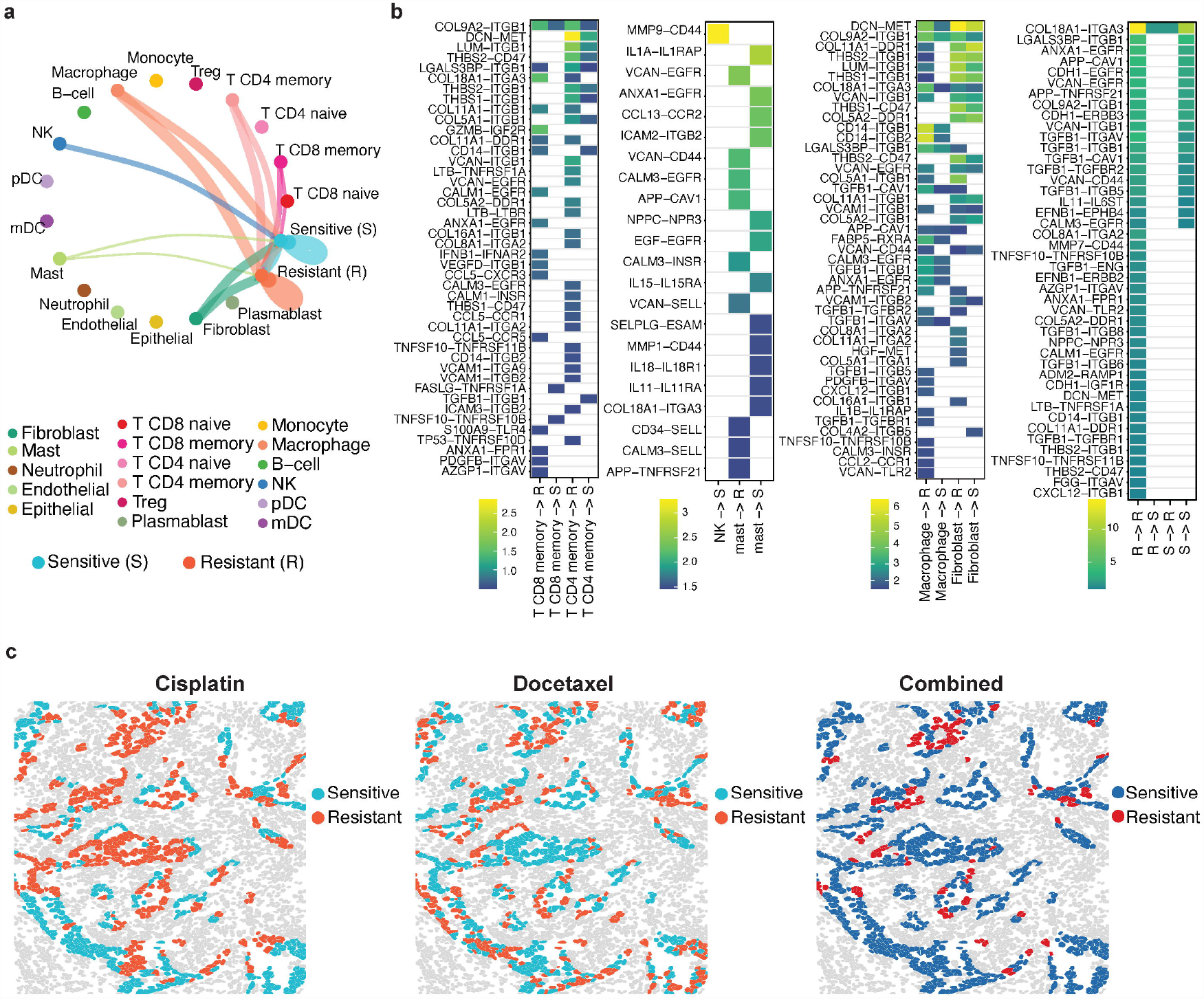
Roles of cell-cell communications in tumor cell resistance. **a**, Summary chord diagram of the identified cell-cell communication network. The chord width is proportional to the interaction strength across different cell types. **b**, Heatmap shows the identified L-R interactions between major cell types and resistant (R) or sensitive (S) cells. **c**, Spatial visualization of cellular response to cisplatin, docetaxel, and their combinations.

The involved L–R pairs that contribute to the cellular crosstalk with tumor cells are further presented in **Fig. 5b**. The gradient colors represent the interaction strength of each L–R pair. Specifically, CD4 and CD8 memory T cells are involved in more L–R interactions with resistant than sensitive cells. *MMP9*–*CD44* is uniquely involved in the interactions between NK and sensitive cells, which is supported by previous study^20,21^. Moreover, fibroblast expressed *DCN* (ligand) shows stronger interactions with resistant tumor cells’ MET (receptor). *DCN* has been reported to interact antagonistically with the *MET* factor (c-Met)^22,23,^ and play roles in cancer development and metastasis^24^. Other L–R interactions including *HGF*^25^–*MET*^26^, *VCAN*^27^–*ITGB1*^28^, and *VCAN*^27^–*CD44*^29^ also play a crucial role in cancer cells forming resistant state against drugs.

SpaRx can be used to explore optimal drug combinations. For example, in addition to cisplatin, SpaRx also identifies the spatially differential cellular response to the other lung cancer drug, docetaxel. As in **Fig. 5c**, some tumor cells that are resistant to one drug appear sensitive to the other drug. Tumor cells sensitive to each of the two drugs are complementary, with Jaccard similarity as 0.381. This result suggests that the combined therapy of cisplatin and docetaxel may overcome resistance and improve therapeutics, which has also been confirmed in clinical trials for patients with unresectable NSCLC^30-32^.

### SpaRx uncovers an orderly pattern of resistant tumor cells

Next we apply SpaRx to the Vizgen MERSCOPE liver cancer SCST data (**Fig. 6a**). In this case, most tumor cells (green colored cells) are confined in three regions with clear boundaries, with some infiltrating tumor cells within the hepatocytes. Specifically, the tumor region on the left (region-1) is surrounded by Kupffer cells, and the other two tumor cell regions (region-2 and region-3) at the right are surrounded by hepatoblasts. SpaRx is applied to predict the tumor cells’ response to a typical liver cancer drug, mitoxantrone. As shown in **Fig. 6b**, SpaRx uncovers tumor cells’ response to mitoxantrone, with both sensitive and resistant cells revealed. Interestingly, sensitive cells mostly present at the outer area, whereas resistant cells majorly locate in the inner area of each tumor region. Such orderly patterns of resistant cells shared by three tumor regions indicate that those resistant cells may share similar molecular characteristics.

**Fig 6:**
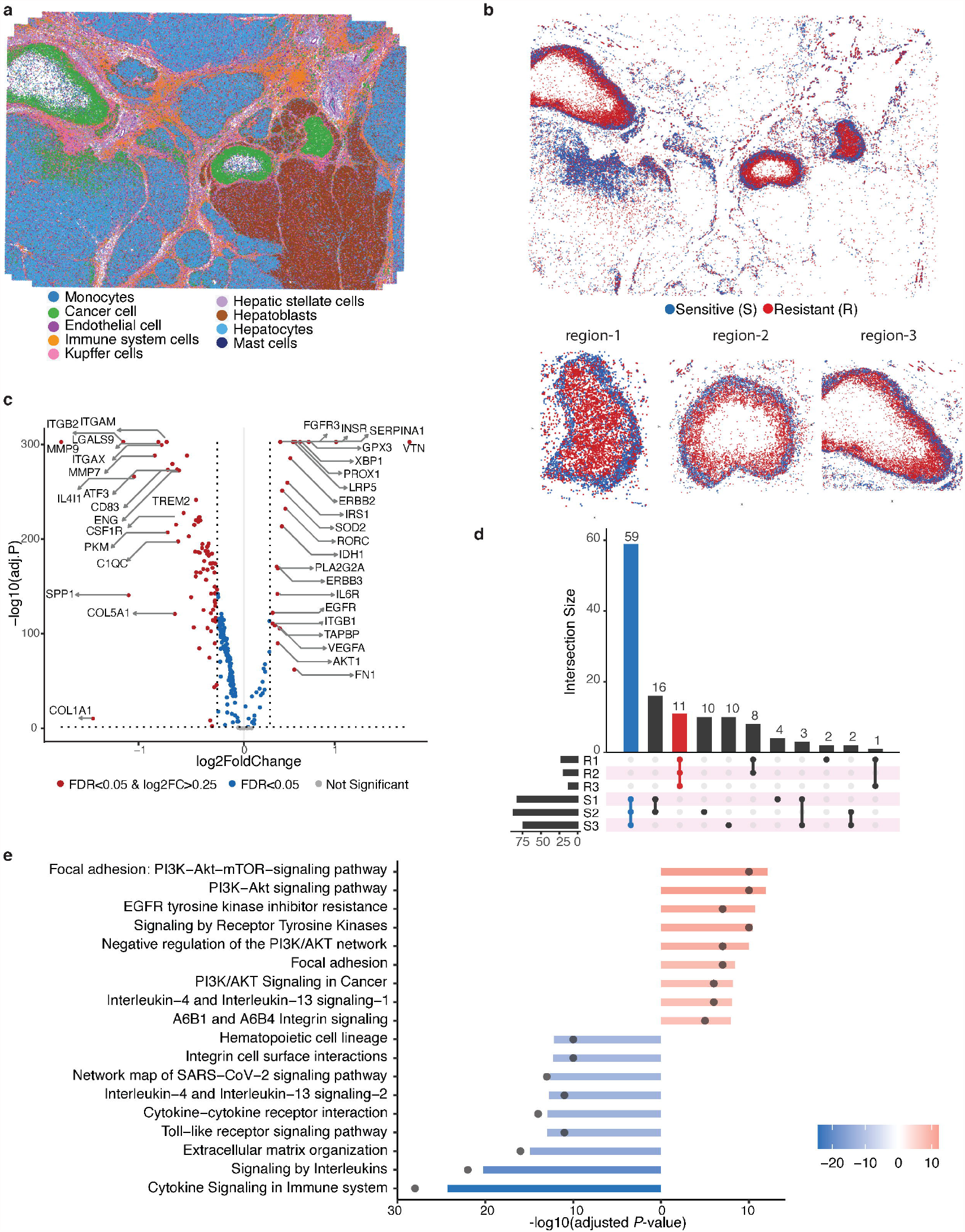
SpaRx reveals drug resistance pattern in liver tumor tissue. **a**, Spatial visualization of the single-cell spatial data from liver tumor tissue. **b**, Spatial visualization of cellular response to mitoxantrone. Red and blue colors denote resistant and sensitive tumor cells to mitoxantrone. **c**, Volcano plot shows the differentially expressed genes (DEGs) of resistant versus sensitive tumor cells in tumor region-1. **d**, Overlaps of DEGs across three tumor regions. **e**, Enriched pathways based on overlapped DEGs across three tumor regions.

To investigate the underlying differences between the resistant and sensitive cells, differentially expressed gene (DEG) analysis is performed for each tumor region. For example, the DEGs of resistant cells at region-1 are shown in **Fig. 6c**, among them *VTN*^33^ and *VEGFA*^34^ are overexpressed. Vitronectin (encoded by *VTN*) has been reported to protect cancer cells from drug-induced apoptosis^33^. *VEGFA* decreases the sensitivity of cancer cells to chemotherapy by suppressing *VEGFA*-mediated autophagy^34^. These over-expressed genes identified in resistant cells may serve as resistance biomarkers and potential therapeutic targets. More importantly, the DEGs of both resistant and sensitive cells across three tumor regions are largely in common (**Fig. 6d**), further confirming that these regions share similar molecular mechanisms for mitoxantrone resistance. Enrichment analysis of these shared DEGs (**Fig. 6e**) among resistant cells (R1, R2, and R3) reveals signaling pathways that are potentially responsible for mitoxantrone resistance, including the focal adhesion-induced PI3K-AKT signaling. In contrast, interleukin signaling and cytokine signaling pathways are enriched in sensitive cells.

## DISCUSSION

The spatial heterogeneity in cells and their microenvironment play critical roles in the treatment of complex diseases such as cancers^35^ and Alzheimer’s diseases (AD)^36^. For example, tumor microenvironment is crucial for tumor cell metastasis^37^ and drug resistance^38^. Recent emerging single-cell spatial technologies utilizing molecular imaging for targeted gene profiling, provide deep insights into the spatial cellular ecosystems^39-41^. These state-of-art technologies help resolve the cellular heterogenous response to drugs, the intercellular communications that contribute to drug resistance, and how tumor ecosystem acquires drug resistance.

In this work, we have developed a novel SpaRx model that leverages the pharmacogenomics knowledgebase with SCST data to systematically reveal spatial complexity of therapeutic response. As to our knowledge, SpaRx is the first method to incorporate the large-scale pharmacogenomics profiles with SCST data, to accurately predict the heterogeneous cellular response to drugs. SpaRx is able to reveal the spatial cell variability in drug response and uncover the underlying biological mechanisms for drug resistance. For example, based on the lung cancer SCST data, we observe the multitude interactions related to tumor cell resistance and identify the spatially adjacent cell interactions that may alter tumor cells’ sensitivity to cisplatin. In addition to cancers, SpaRx also holds the promises for repurposing anti-cancer drugs for complex diseases such as AD, which is also known for its complexity and heterogeneity. Collectively, SpaRx is anticipated to reveal the mechanisms of drug resistance, prioritize tailored drugs for complex diseases, and provide clues for drug repositioning.

Given the advantages of SpaRx, there are several aspects that SpaRx can be improved. First, current single-cell spatial technologies are still not able to detect sufficient number of genes, which may limit the potentials of SpaRx in some degree. Future advances in spatial technologies that captures more genes and less dropouts will help enhance the SpaRx model. Second, with the rapid development of single-cell spatial omics technologies^42^, SpaRx can also be improved through incorporating spatial multi-omics. Though the current version of SpaRx that enables the predictions of drug responses based on SCST data, SpaRx can be improved by utilizing new data types, e.g., single-cell spatial ATAC-seq profiles^43^, thus to further unveil the underlying mechanisms such as the upstream cis-regulatory elements and associated transcription factors involved in drug resistance.

## MATERIALS AND METHODS

### Data sources and preparation

1. The GDSC^10^ and CCLE^9^ cell-line-based drug screening database. The gene expressions of cell lines and the sensitivity profile (IC50) of drugs are downloaded from the GDSC and CCLE database. The binary drug responses for each cell line are obtained from previous studies^11,13^. Here for GDSC and CCLE database, cell lines and drug information without missing values are retained and integrated based on overlapped drug compounds. Collectively, we obtained 1,280 cell lines with drug response information across 80 shared drug compounds.
2. The NanoString CosMx lung-13 SCST data^5^ and the Vizgen MERSCOPE liver cancer-1 SCST data^6^.

### Benchmarking data

The benchmarking datasets are generated from the collected 1,280 cell lines. Here we randomly select a proportion of cell lines (*p*) as the source domain, whereas the remaining cell lines (1-*p*) are used to synthesize the target domain. For the target domain, the gene expression profiles of the remaining cell lines (1-*p*) are further mixed randomly to mimic tumor cell complexity. These mixed data is then downsampled to assure the total counts are comparable to the single-cell level, which allows the synthesized gene expression data in the target domain mimic that of single tumor cells in SCST data. Specifically, to mimic the complexity of tumor cells, we select two to ten cell lines from the remaining cell lines (1-*p*), then combine their transcriptomic profiles as one tumor cell profile. In order to better mimic real tumor cell, if the total counts of the resulting tumor data exceed 2,000, we downsample it accordingly. In this way, the synthesized gene expression data serving as the target domain is more likely to resemble the real heterogenous tumor cells in SCST data.

Moreover, four benchmarking scenarios are included. 1) Different numbers of cell lines in the source data. We choose different proportions of cell lines as the source domain, i.e., *p* = 10%, 30%, 50%, respectively. The remaining cell lines (1 − *p*) are used to generate the target domain. 2) Different levels of noises. For both source and target domain based on the setting of *p* = 30%, we add extra noises randomly sampled from normal distributions *N*(0, *σ*^2^), with standard deviation *σ* as 1, 1.5, and 2, respectively. 3) Different levels of dropouts. For both source and target domain based on the setting of *p* = 30%, dropouts are simulated by replacing the gene expression values with zeros, to ensure the proportions of zeros among all gene expression values are 70%, 80%, and 90%, respectively. 4) Different numbers of genes. Based on the setting of *p* = 30%, the number of genes in the cell line profiles is reduced to 2k, 853, and 500 genes. The 2k genes are randomly selected, while the 853 and 500 genes are selected based on the RNA panels used in the NanoString CosMx and the Vizgen MERSCOPE data, respectively.

### SpaRx model

#### Source domain

The GDSC and the CCLE data are used as the source domain, denoted as 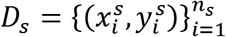. Here *s* represents the source domain (the cell-line based drug response profiles), 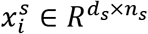 represents the gene expression, with *i* representing a cell line, *d*_*s*_ representing the number of genes, and *n*_*s*_ denoting the number of cell lines. The cell-line similarity graph *A*_*s*_ is constructed using mutual nearest neighbors (MNN)^44^, with the number of mutual nearest neighbor as *k*. In the graph *A*_*s*_, each node *v*_*i*_ represents a cell line *i*, and if two nodes *v*_*i*_ and *v*_*j*_ are connected, it means that the corresponding gene expression profiles 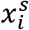 and 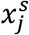 are similar.

#### Target domain

The SCST data is used as the target domain. Each of the SCST data is represented by 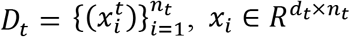, where *t* denotes the target domain (the SCST data), *d*_t_ denotes the number of genes, *n*_t_ represents the number of cells, and *i* represents a cell in the SCST data. A spatial cell graph *A*_*t*_ is constructed according to cell locations using *k* -nearest neighbors. If two cells *i* and *j* are spatially adjacent, then the corresponding nodes *v*_*i*_ and *v*_*j*_ are connected in *A*_*t*_.

The SpaRx model uses cell lines in the source domain and cells in the target domain as samples, gene expressions as features, and drug responses as outcomes. SpaRx is composed of three components: 1) feature extractor to extract gene expression features from the source and the target domain, 2) drug response predictor for both cell lines and single cells, and 3) global and drug-specific discriminators. The final output of the SpaRx is the predicted drug responses of each cell in the target SCST domain.

##### 1) Feature Extractor

The shared feature extractor *T*_*r*_ is composed of multi-head graph transformer^15^ layers to project the graph representation of the cellular transcriptomics data to a latent space in which cells that demonstrate similar responses to treatments are close to each other. Briefly, for a cell line *i* from the GDSC or the CCLE data or a cell *i* from the SCST data, the propagation of the graph transformer from the *l* layer to the *l* + 1 layer is defined as:

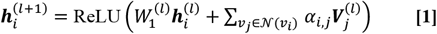

where *i* represents either a cell line 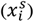 in the source domain or a cell 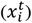 in the target domain, *j* represents a neighbor cell line or cell in their corresponding graph (i.e., *v*_*j*_ ∈ 𝒩(*v*_*i*_)), the rectified linear unit (ReLU)^45^ is used as the nonlinear gated activation function. When *l* = 0, 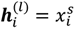 for the cell line data and 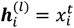 for the SCST data. The attention module is defined as: 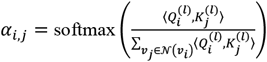, where:

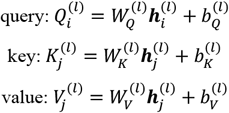

and 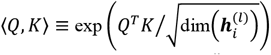. The multi-head attentions are concatenated. In this way, we obtain the latent representation, 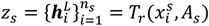, as the extracted features for source domain, and 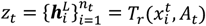 as the extract features for target domain, respectively. The feature extractor *T*_*r*_ is shared by both source and target domain.

##### 2) Drug response predictor

The predictor (*F*_*y*_), a fully connected classifier, is designed to classify the drug response results using latent features from the feature extractor *T*_*r*_. It is trained by minimizing the differences between the predicted source labels and the source domain labels of drug response (ground truth labels) by the cross-entropy loss, which is formulated as:

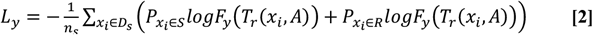

where 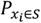 is the probability of *x*_*i*_ belonging to drug sensitive, 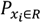 is the probability of *x*_*i*_ belonging to drug resistance. *F*_*y*_ is the response predictor and *T*_*r*_ is the feature extractor. *A* represents the graph *A*_*s*_ in source domain. Here the drug response predictor is shared by both source and target domain.

##### 3) Discriminators

A global discriminator *G*_*d*_ is trained to align the latent representations of source and target domain. Here the loss of the global discriminator is formulated as:

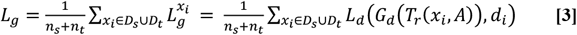

where *L*_*d*_ denotes the cross-entropy, *G*_*d*_ denotes the global discriminator, *T*_*r*_ is the feature extractor, and *d*_*i*_ is the domain label for the input *x*_*i*_(*d*_*i*_ = 0 for source domain, *d*_*i*_ = 1 for target domain). *A* represents the graph *A*_*s*_ when *x*_*i*_ ∈ *D*_*s*_ and *A*_*t*_ when *x*_*i*_ ∈ *D*_*t*_.

Drug-specific discriminators 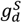 and 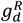 : 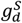 and 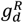 are used to match the latent representations from source and target domains under drug-sensitive and drug-resistant category, respectively. Both drug-specific discriminators are trained to minimize the differences in the latent representations of source and target domain under each drug category. The output of the drug response predictor (*F*_*y*_(*T*_*r*_(*x*_*i*_))) is used to show the probability of being included into each drug category. The loss for each discriminator is calculated using cross-entropy:

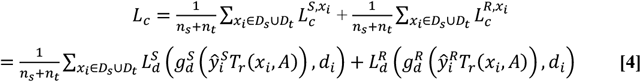

where 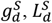 and 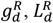 are the drug-specific discriminator loss and its cross-entropy loss associated with drug categories, respectively. 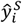 and 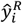 is the predicted probability of the input *x*_*i*_ belonging to drug-sensitive or drug-resistant category, i.e., *F*_*y*_(*T*_*r*_(*x*_*i*_)). *d*_*i*_ is the label for the input *x*_*i*_(*d*_*i*_ = 0 for source domain, *d*_*i*_ = 1 for target domain). *A* represents the graph *A*_*s*_ when *x*_*i*_ ∈ *D*_*s*_ and *A*_*t*_ when *x*_*i*_ ∈ *D*_*t*_.

#### Loss function

Given the three major components, feature extractors, domain discriminators, and label classifier in our model, the final learning objective is formulated as:

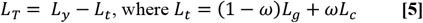

In the loss function, ω is a dynamic adversarial factor which balances the relative weight of the global and the drug-specific discriminator loss. During the training, ω is dynamically updated according to the losses of the three discriminators: 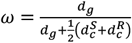, where, 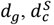, and 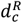 are the proxy 𝒜-distances^46^ between the source and the target domains for the three domain discriminators, respectively. Specifically, for the discriminator, 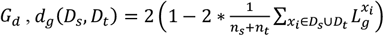; for the discriminator 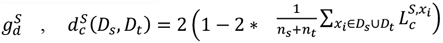, ; and for the discriminator 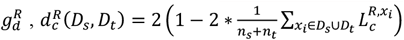, where *n*_*s*_ represents the number of cell lines in source domain, and *n*_*t*_ is the number of cells in target domain.

The SpaRx model is trained using the stochastic gradient descent (SGD) optimizer. In the SpaRx model, the parameters including the number of adjacent neighbors or mutual nearest neighbors in graph construction, as well as the latent dimensions in graph transformer layers, are determined through grid-based hyper-parameter fine tuning. The hyperparameters used in the final model are: for the SGD optimizer, momentum=0.9 and weight decay=5*e*^−5^; the learning rate is set to 1*e*^−3^; gradient clip threshold at 5; the number of graph transformer layers is 2, with the dimensions of 512 and 64, respectively. After the model training, SpaRx accurately predicts the drug response labels of cells in the spatial data and uncovers the spatially heterogeneous responses to different types of drugs.

### Benchmarking methods and comparison measurement

To evaluate the performance of SpaRx, we compare it with four deep learning models, including SpaRx-GAT, SpaRx-GCN, scDEAL^13^, SCAD^14^, and four machine learning methods including random forest (RF), support vector machines (SVM), lightGBM and XGBoost. SpaRx-GAT is built based on the SpaRx model, with the feature extractor as GAT^16^ layers, rather than the graph transformer^15^. SpaRx-GCN uses GCN^17^ layers as the feature extractor. scDEAL^13^ and SCAD^14^ are proposed to predict single cell response to cancer drugs by integrating large-scale bulk cell-line data and scRNA-seq data. To evaluate the performance of each model, we use the F1 score to assess the agreement between the predicted drug response and the ground truth. The F1 ranges from 0 to 1 refering to the increasing match between the predicted drug response with ground truth. With *TP* denoting true positive, *FP* representing false positive, and *FN* representing false negative, F1 score is calculated by 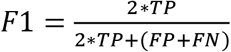. Additional metrics including precision, recall, AUROC, and AUPR are included for comprehensive evaluation of SpaRx and benchmarking methods.

### Identifying surrounding microenvironment, L-R interactions, and adjacent cell communications

When characterizing the surrounding microenvironment, we have summed up the adjacent (with 5 nearest neighbors) cells (by each cell type) around resistant/sensitive cells. After dividing the total number of cells within each cell type, we obtain the percentage of different cell types in the microenvironment of resistant/sensitive cells. To identify L-R interactions, our previous tool spaCI^19^ is used here for SCST data. With the L-R interactions, we further characterize the adjacent cell communications with interaction strength. Specifically, for an L-R interaction pair, we define its interaction strength as the multiplication of their average expression values among adjacent cells, where the top and bottom 10% expressions of the ligand and the receptor are ignored. The interaction strength of all identified L-R pairs is then summarized as the interaction strength between two cell types. Thus, the higher value of the interaction strength, the stronger the two cell types adjacently interact.

## DATA AVAILABILITY

**NanoString CosMx SMI data:** The single-cell spatial dataset (Lung-13), profiled by CosMx SMI on Formalin-Fixed Paraffin-Embedded (FFPE) samples of the non-small-cell lung cancer (NSCLC) tissue^5^, is available from https://nanostring.com/products/cosmx-spatial-molecular-imager/ffpe-dataset/. **Vizgen MERSCOPE data:** We includes the Vizgen MERFISH liver cancer 1 dataset that contains a MERFISH measurement of a 500 gene panel. Data is downloaded from https://info.vizgen.com/merscope-ffpe-solution, which includes the list of detected transcripts, gene counts per cell matrix, and additional spatial cell metadata. The gene expression profiles of GDSC and CCLE cell lines are downloaded from https://www.cancerrxgene.org/ and https://depmap.org/portal/. The gene expression profile data of TCGA lung cancer patients, including lung adenocarcinoma and lung squamous cell carcinoma patients, are downloaded from the UCSC Xena database (http://xena.ucsc.edu/). The corresponding response information to cisplatin is retrieved from previous studies^47^, where responders (including complete response and partial response) and non-responders (including stable disease and progressive disease), are characterized according to the RECIST standard^48^.

## CODE AVAILABILITY

SpaRx is provided as a Python package available at https://github.com/QSong-github/SpaRx, with detailed functions for the general applicability on different SCST data.

## COMPETING INTERESTS

The authors declare no competing interests.

## FUNDING

QS is supported in part by the Bioinformatics Shared Resources under the NCI Cancer Center Support Grant to the Comprehensive Cancer Center of Wake Forest University Health Sciences (P30CA012197). QS is also supported by the American Cancer Society Institutional Research Grant Pilot. JS is partially financially supported by the Indiana University Precision Health Initiative and the Indiana University Melvin and Bren Simon Comprehensive Cancer Center Support Grant from the National Cancer Institute (P30CA082709). JS is also supported by R01LM013771.

## MATERIALS & CORRESPONDENCE

Correspondence and requests for materials should be addressed to QS or JS.

## FIGURE LEGENDS

**Supplementary Fig. 1: Performance of SpaRx and additional benchmarking methods. a**, Performance of SpaRx and other methods (SCAD, scDEAL, LightGBM, and XGBoost) are measured by the F1 scores across 80 different drug compounds. Each point represents the F1 score of SpaRx versus an alternative method on one type of drug. **b**, Boxplot of F1 scores in different scenarios, including different source data sizes, noise levels, dropout levels, and number of genes.

**Supplementary Fig. 2: AUROC scores of benchmarking methods in different scenarios**.

**Supplementary Fig. 3: AUPR scores of benchmarking methods in different scenarios**.

**Supplementary Fig. 4: Precision values of benchmarking methods in different scenarios**.

**Supplementary Fig. 5: Recall values of benchmarking methods in different scenarios**.

**Supplementary File 1: Comprehensive evaluation of SpaRx with benchmarking methods**.

## Notes

### Competing Interest Statement

The authors have declared no competing interest.

